# Regulation by HSP70/90 in the different tissues and testis development of male cattle (Cattle-yak and Yak)

**DOI:** 10.1101/393371

**Authors:** Penggang Liu, Sijiu Yu, Yan Cui, Junfeng He, Qian Zhang, Jun Liu, Liangli Song, Yuanfang Xu, Ting Wang, Shengnan Zou, Hui Li

## Abstract

HSP70/90 play important role in testis develop and spermatozoa regulation, but the contact of HSP70/90 with infertility in cattle is unclear. Here, we focus on male cattle-yak and yak, which to investigate the expression and localization of HSP70/90 in different tissues, and explore the influence of HSP70/90 to infertility. In our study, a total of 54 cattle (24 cattle-yak and 30 yak) were examined. The HSP90 mRNA of cattle-yak was cloned first and found amino acid variation in HSP90, which led to difference at protein spatial structure compare with yak. To investigate whether the expression of HSP70/90 mRNA and protein are different in cattle-yak and yak, we used real-time quantitative PCR (qRT-PCR) and Western blot (WB) to examine them. We found that the expression level of HSP70/90 mRNA and protein are disparity in different tissues and testis development stages, and obviously high expression was observed in testicle during juvenile and adult, Moreover, it‘s interestingly in which the HSP70 expression is significant high in yak whereas HSP90 in cattle-yak (P<0.01). On this bases, we detect the location of HSP70/90 in testis by immunohistochemical (IHC) and immunofluorescence (IF), the results demonstrate that HSP70/90 were located in the epithelial cells, spermatogenic cells and mesenchymal cells. In summary, our study proved the expression of HSP70/90 are different in tissues, and the expression of HSP90 is obviously high in testis of cattle-yak, which propose that the infertility of cattle-yak may cause from up-regulating of HSP90.

## Introduction

Cattle-yak (*Bos* cattle-yak) and yak *(Bos grunniens)* are important animals for the local herdsmen in the Qinghai-Tibet Plateau in China. Cattle-yak as a hybrid exhibit obvious heterosis of yak and cattle, such as tall variety, robustness, fast growth speed, drought tolerance, and disease resistance (Wiener et al. 2003). Very strong resistance to cold and high altitude is necessary for this species to adapt to the cold environment. Heat shock protein 70/90(HSP70/90) plays key roles in protecting the body to avoid irritation. In this study, we determined the relationship between HSP70/90 and tissue specificities of cattle and the regulation of testicular development and sperm production.

Recent studies have shown that HSP70/90 could protect cells from damage, organ failure, and body shock, among others. HSP70/90 not only directly or indirectly participates in the tumorigenesis, autoimmunity, wound healing, cell migration and transshipment for growth and development but also protects reproductive cell activity and regulates protein metabolism of organisms (Carper et al. 1987). Previous studies demonstrated high concentrations of HSPs in developing tissues and tumor cells, such as in stomach cancer, colon cancer, liver cancer, breast cancer, and lung cancer (Manjili et al. 2002; Weng et al. (2014) showed that the HSPA2 expression levels significantly differed in the various tissues of male yak, especially in the testes, followed by the brain, kidney, heart, lung, and liver, and its weakest expression was found in the spleen (Weng et al. 2014). That showed HSP70/90 play important role in tissue specificities of female yaks as well as lactating function (Liu et al. 2017). This study was performed to investigate whether HSP70/90 was differentially expressed in the various tissues of cattle-yak and yak.

The testis is one of the most important male reproductive organs, and the place of its cell proliferation and development is important. Many factors are responsible for the infertility of male cattle-yak. These factors include testicular dysplasia, hindered spermatogenesis, metamorphosis phase, and sustentacular cells assisting the germ cell differentiation in the seminiferous tubule (Lu and Zi 2014). Thus, studies on the fertility of male cattle-yak and male yak have focused on distinguishing histologic structural development, the effect of gene expression genes, and the intensity of protein involved in the production and control of sperms (Shao et al. 2016).

Cattle-yak infertility is not only reflected histologically, but studies also showed that the differential expression of related gene and protein generated by sperms is another factor for infertility. HSP is mainly involved in biological processes, such as sperm movement, energy metabolism, protein processing, and oxidative stress. Asthenozoospermia creatures have low content of eight differential proteins containing triosephosphate isomerase, glycerol kinase, and HSP (Siva et al. 2010; Sönmez et al. 2017). HSP70 plays an important role in sperm meiosis, sperm maturation, and sperm–egg recognition. In the sperm of human beings, the expression of HSPA2 in the elongated sperm and the sperm cells is significantly higher than that of the primary spermatocyte (Motiei et al. 2013). In addition, temperature plays an important role in spermatogenesis. Hyperthermy induce cells to express HSP, which improves heat tolerance, protects protein structure from injuries, and repairs the injury of the protein structure and functional integrity. High temperature triggers cell apoptosis and causes the redistribution of HSP in the cells. For example, the cytoplasm moves to the nucleus and nuclear membrane to protect cells from damage. Different degrees of damage result in diverse consequences. Several damages stop cell proliferation and activate cell repair mechanisms simultaneously, while some damages directly initiate cell death (Chen et al. 2013). Heating the testis for 10 min at 43 °C does not evidently change the morphology of the seminiferous tubule. Spermatocyte and circular sperm cells are sensitive to heat shock, and apoptosis occurs with heating (LI et al. 2016). Sudden change in temperature during animal breeding stage can cause HSP60/70/90 to rapidly respond and become highly expressed in the gene level to improve the ability to protect gap-associated protein from disturbance (Mikulski et al. 2011). However, it is still a blank that the correlation between male sterile and HSP70/90 of mules and cattle-yak. Therefore, this study will provide theoretical basis for hybrid sterility.

## Results

### 1. Analysis physical and chemical properties of HSP90

The cDNA sequence of HSP90 was cloned and submitted to Genbank with accession number KF690731.

The nucleotide sequence and predicted amino acid sequence of HSP90 were presented. HSP90 nucleic acids were 3064 bp long. Results from the analysis of the contig showed that the predicted HSP90 cDNA of 3064 bp contained an ORF of 2202 bp from 146 bp to 2347 bp. The small three ORFs (ORF1, ORF2, and ORF3) coded for a nonfunctional protein was composed of 123 and 171 amino acids. Analysis of the HSP90 cDNA sequence in our study confirmed this conclusion. To verify this result, three primers H90-1, H90-2, and H90-3 were used to clone the full-length ORF of HSP90. RT-PCR results show the successful isolation of a cDNA fragment of 2202 bp from the cattle-yak total RNA of the heart. Finally, this confirmed cDNA sequence was deposited in the GenBank under accession number KF690731. To obtain the genomic DNA of HSP90, the publicly available cow genome database at the NCBI Bovine Genome Resources (http://www.ncbi.nlm.nih.gov/projects/genome/guide/cow/) was screened using the HSP90 cDNA sequence as a query. A cow *(Bos taurus)* contig (GenBank Accession No. NM_001012670) which encompasses the entire HSP90 gene was identified by BLASTGen analysis.

The basic physical and chemical properties of HSP90 were analyzed. The atomic number of the protein coding region is 11710. HSP90 has a molecular formula of C_3733_H_5950_N_986_O_1202_S_27_, molecular weight of 847408.84 Da, and theoretical isoelectric point of 4.841. HSP90 has a half-life of approximately 30 h, instability index of 42.77, fat soluble index of 79.80, and average hydrophobicity index of -0.744.

A molecular phylogenetic tree was constructed to analyze the evolutionary relationship of HSP90 nucleotide sequences (Fig.1).

**Fig 1.**
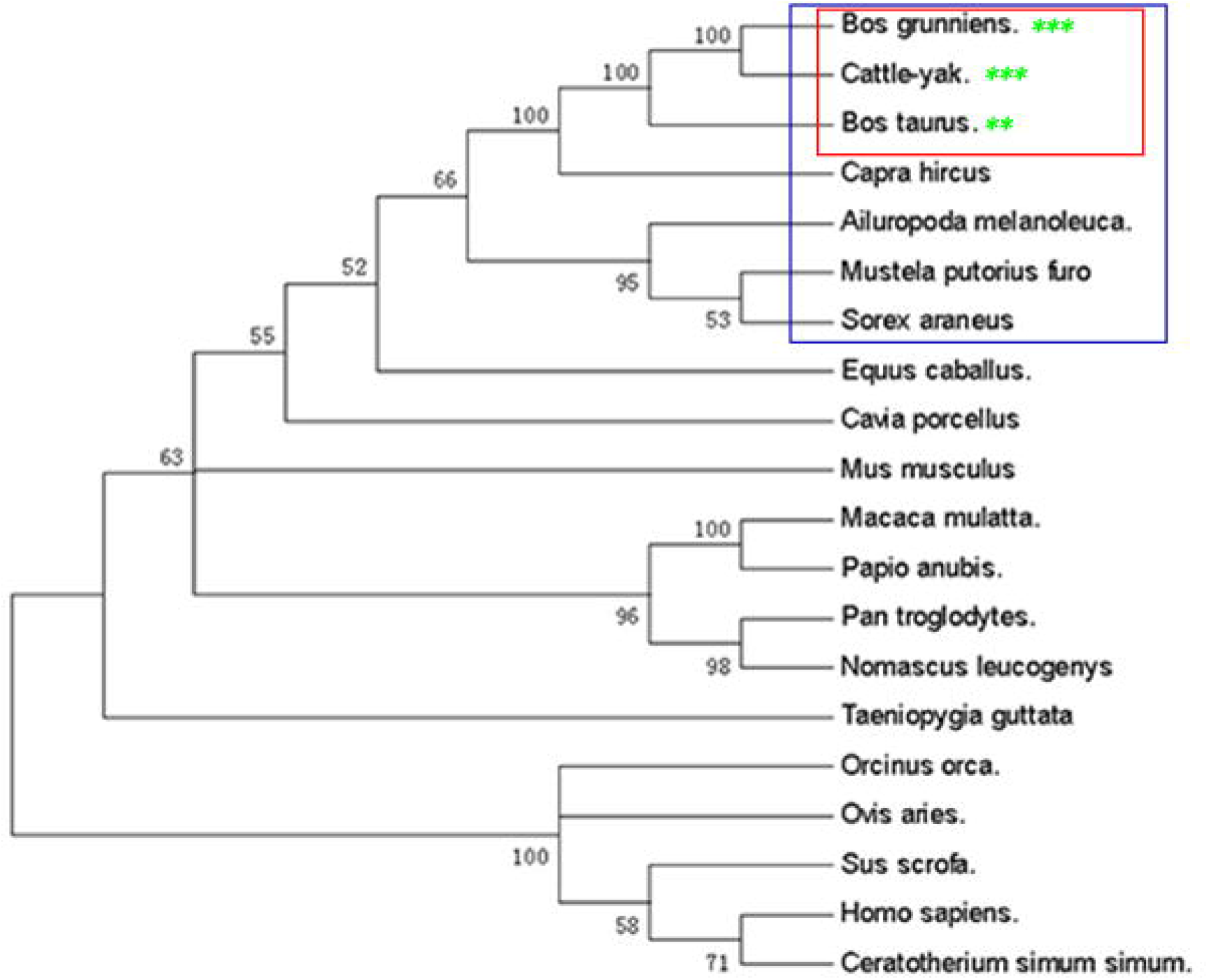
The HSP90 phylogenetic tree of cattle-yak. The phylogenetic tree was constructed with CLC main workbench software using the neighbor-joining method for the amino acid sequences of HSP90 from the Swiss-Prot databank/GenBank.

Analysis of the gene family tree showed that the cattle-yak HSP90 evolutionarily shared a higher sequence identity with *B. grunniens, B. taurus, Capra hircus, Ailuropoda melanoleuca, Mustela putorius furo, Sorexaraneus, Equus caballus*, and *Cavia porcellus*.

### 2. Analysis structure specificity with proteins of HSP90

The predicted amino acid sequence of HSP90 was aligned with known *B. grunniens* sequences through BLASTp. The cattle-yak HSP90 protein sequence shared a low percentage of similarity to other known HSP90 protein sequences. This result indicated that the HSP90 proteins were different from other members in the HSP family. The amino acid sequence of HSP90 was approximately 99.56%, 99.33%, 96.91%, 87.83%, and 86.90% identical to those of HSPs from *B. grunniens, B. taurus, C. hircus, E. caballus*, and *Mus musculus*. The highest level of similarity appeared near the C-terminal, while the very low similarity was found in the N-terminal and in the middle of the amino acid sequence.

The cattle-yak HSP90 protein sequence showed that the mutational nucleotide caused the change of amino acids, such as the GLY into GLU, FGI into LEF, GLN into ARG, and MET into ILE (Fig. 2). The most important result is the change in the protein spatial structure and differences in protein function, such as the H-key number (397), spiral number (30), link number (23) and corner number(89). In addition, compare with yak, the initiating terminal and terminal of the cattle-yak are longer. The results from the SDS-PAGE analysis indicated that the fusion protein HSP70 has a molecular weight of approximately 70 kDa, whereas the fusion protein HSP90 was approximately 84.7 kDa.

**Fig 2.**
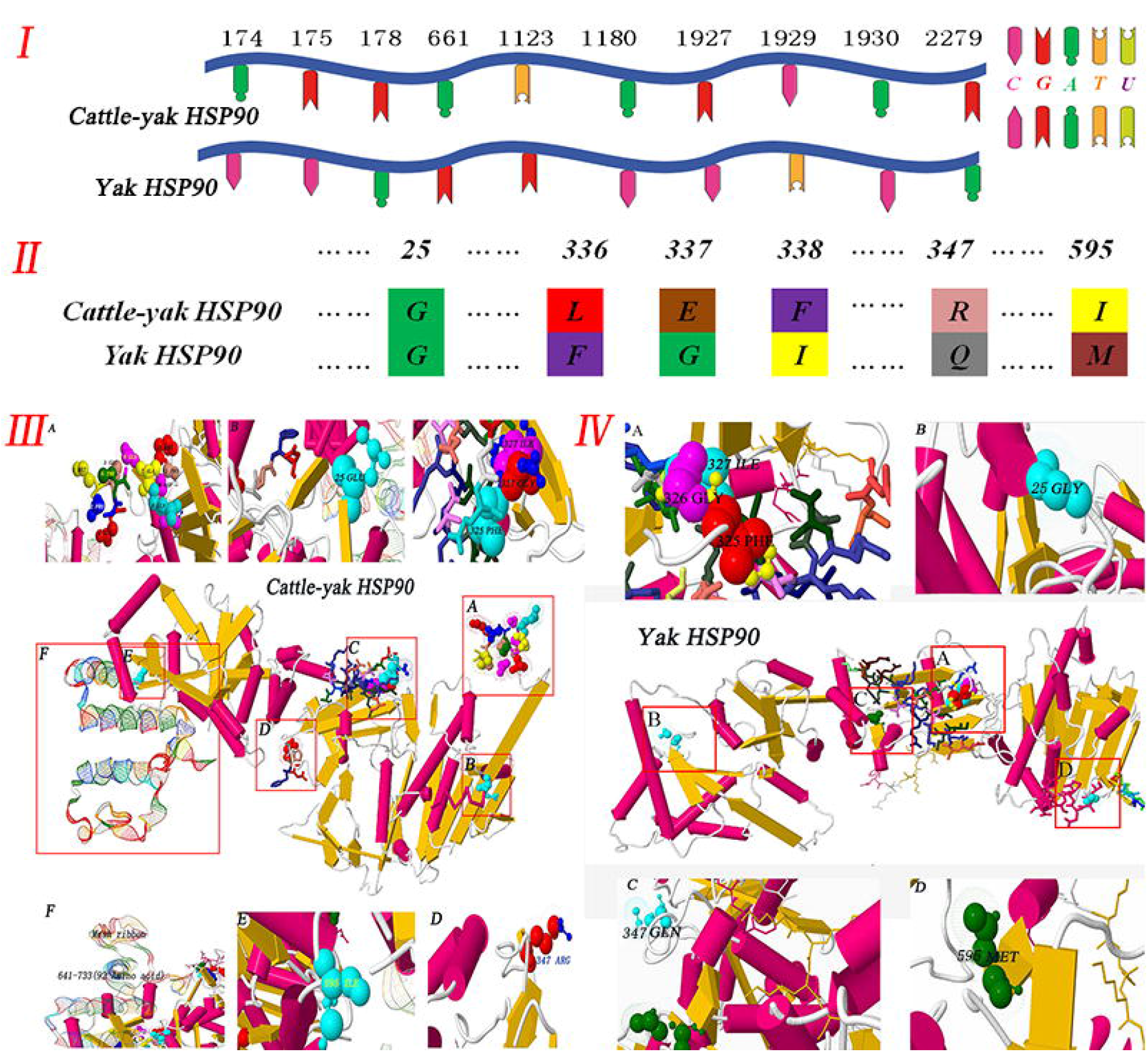
Analysis of HSP90 protein structure of cattle-yak and yak. Iand☐. The HSP90 nucleotide and amino acid sequence analysis that the sites and types of mutation in cattle-yak and yak. III. The HSP90 protein structure of cattle-yak. (A) The initiating terminal of the cattle-yak is much more than that of yaks, as follows: MET, PRO, GLU, GLU, THR, GLN, ALA, GLN, ASP, PRO, PRO. (B) (D), (E) Mutation site and amino acids are as follows: 25(GLU), 36(THR), 37(PHE), 38(TYR), 347(ARG), 595(ILE). (C) More than 92 amino acids at the terminal. IV. The HSP90 protein structure of yak. (A), (B), (C), (D) Mutation site and amino acids are as follows: 14(GLY), 25(THR), 26(PHE),27(TYR), 336(GLN), 584(MET).

### 3. Expression and distribution of HSP70/90 in the non-reproductive system

The HSP70/90 mRNA in the tissues of the cattle-yak and yak were investigated through real-time RT-PCR (Fig. 3-I), with the total RNA isolated from cattle-yak and yak tissues as a template. As shown in Fig. 3-II, we examined six tissues in cattle-yak and yak, the expression of HSP70/90 are widely distributed in all of them. According from our data, HSP70/90 are both reduce in cattle-yak, the trend from lung, cerebellum, kidney, liver, heart to spleen. There is no significant difference about HSP70 in these tissue, but interestingly the expression levels of HSP90 were consistently higher than HSP70 in almost all the tested tissues (P<0.01). At the meantime, although the expression of HSP70/90 are both higher in cerebellum, kidney and heart in yak, the expression of HSP70/90 in different tissues and level are irregular. Taken together our data suggest that the expression of HSP90 was consistently higher than HSP70 in all the tested tissues (P<0.01) in cattle-yak but irregular in yak.

**Fig 3.**
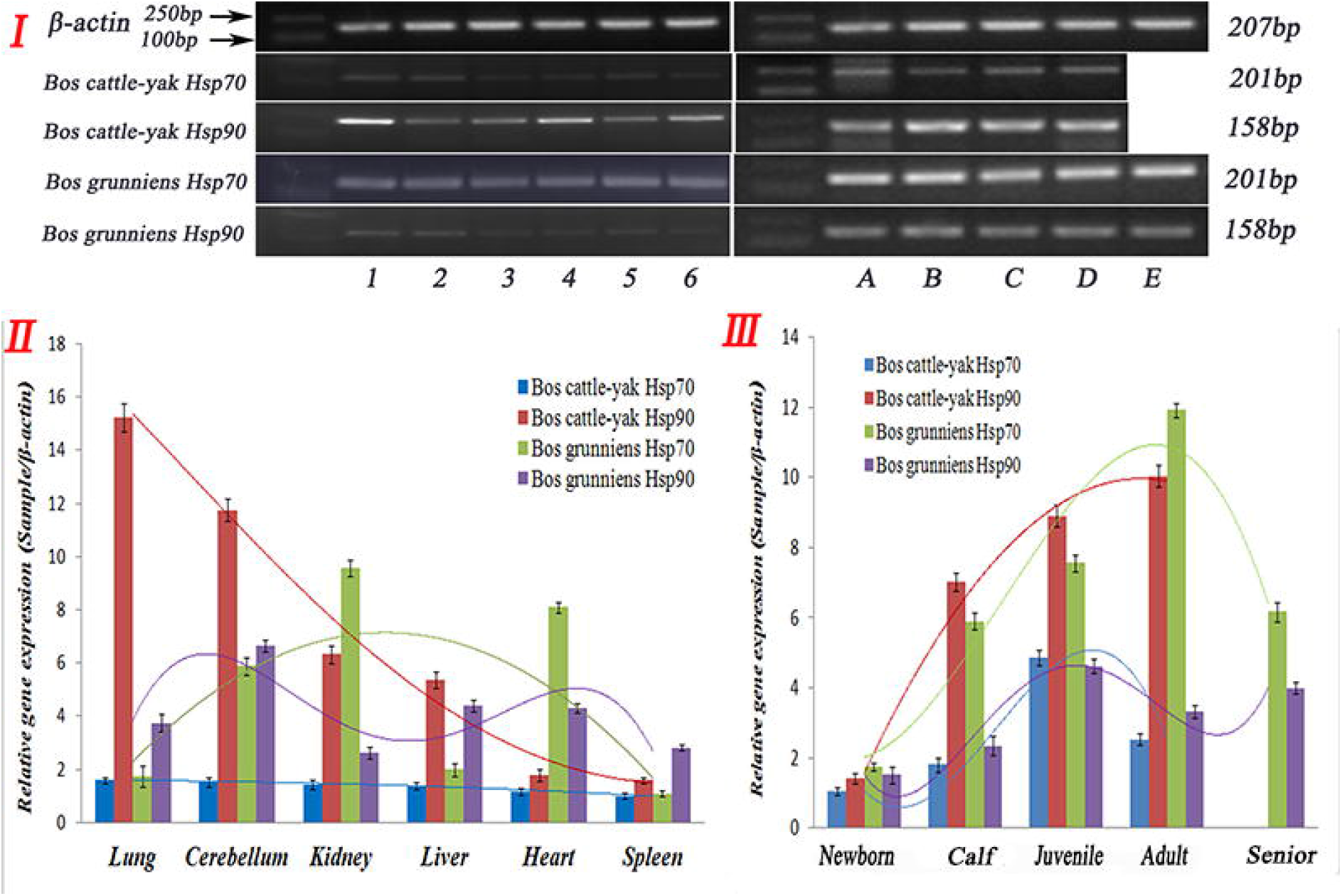
The gene expression in different tissues of cattle-yak and yak. ☐. The results of RT-PCR. (1) Lung; (2) Cerebellum; (3) Kidney; (4) Liver; (5) Heart; (6) Spleen. (A) Newborn; (B) Childhood; (C) Juvenile; (D) Adult; and (E) Senile. II. The gene expression levels of HSP70 and HSP90 in non-reproductive system (cattle-yak and yak) III. The gene expression levels of HSP70 and HSP90 in testicular tissue of male cattle (cattle-yak and yak). Note. The curved lines represent the expression trend of HSP70/90 protein in different organizations and testis development stages. Different colors represent different expressions of HSP70/90 in the organizations and testicular tissue of cattle-yak and yak.

The HSP70/90 was mainly obversed in the kidney tubules, cardiac muscle cells, hepatocytes, Purkinje cells, and cerebellar medulla (Fig. 4-1 and II). Positioning analysis indicated that the HSP70/90 protein was mainly concentrated in the connective tissue and epithelia. Thus, the proteins were mainly concentrated in the cell membrane and in the cytoplasm but not in the nucleus. Furthermore, the protein expression levels of HSP90 were markedly lower than those of HSP70 in almost all the tested tissues (P<0.01), except in the lung (Fig. 4-II I).

**Fig 4.**
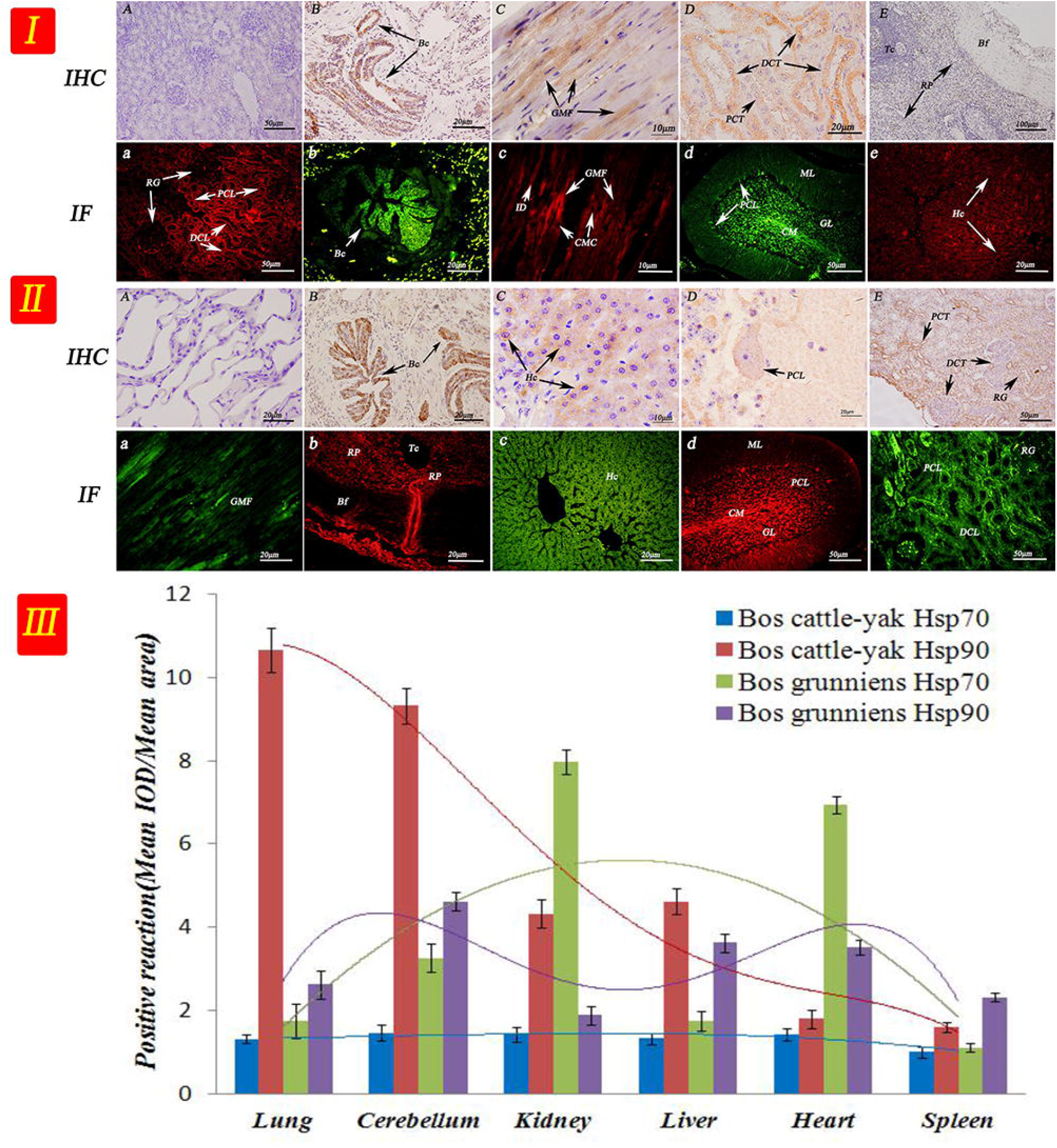
IHC and IF localization of HSP70/90 in the non-reproductive system. ☐. (A) The control sections collected from the kidneys of yak, without immunoreactions (negative control). (B), (D) Positive staining for HSP90 was observed in the terminal bronchioles, distal convoluted tubule and proximal convoluted tubule of the lungs and kidney. (C), (E) Positive staining for HSP70 was observed in the cardiac muscle fibers and red pulp of heart and spleen. (a), (c), (e) Positive staining for HSP70 was observed in the distal convoluted tubule, proximal convoluted tubule, cardiac muscle fibers and hepatocytes in kidney, heart and liver (b), (d) Positive staining for HSP90 was observed in the terminal bronchioles, cerebellar medulla, granular layer and purkinje cell layer of the lungs and cerebellum. . (A) The control sections collected from the lung of yak, without immunoreactions (negative control). (B), (D) Positive staining for HSP70 was observed in the terminal bronchioles, cerebellar medulla, granular layer and purkinje cell layer of the lungs and cerebellum. (C), (E) Positive staining for HSP90 was observed in hepatocytes, distal convoluted tubule and proximal convoluted tubule of the liver and kidney. (a), (c), (e) Positive staining for HSP90 was observed in the cardiac muscle fibers, hepatocytes, distal convoluted tubule and proximal convoluted tubule of heart, liver and kidney. (b), (d) Positive staining for HSP70 was observed in the red pulp, cerebellar medulla, granular layer and purkinje cell layer of the spleen and cerebellum. Note. Terminal bronchiole (TB); hepatocyte (Hc); central vein (CV); cardiac muscle fibers (CMF); distal convoluted tubule (DCT); Purkinje cell layer (PCL); renal glomerulus (RG); biofilm (Bf); trabecula (Tc); red pulp (RP); white pulp (WP); cerebellar medulla (CM); molecular layer (ML); granular layer (GL); and proximal convoluted tubule (PCT). The result of optical density analysis value. The curved lines represent the expression trend of HSP protein in different organizations, and different color represents different proteins.

### 4. Expression and distribution of HSP70/90 at the different development stages of the testis tissue

As we found the difference of HSP70/90 between cattle-yak and yak in six non-reproductive tissue above, we examined testis further. As shown in Fig. 3-III, HSP70/90 gene expression levels showed significant differences in the cattle-yak and yak testicular tissue at different development stages (P<0.01). In cattle-yak, the expression of HSP70 showed obviously tend from newborn to adult, it increased to highest in juvenile and then tend to decreased, at the same time, HSP90 keep increasing until adult. It is worth notice that the expression of HSP90 was significant higher than HSP70 in every develop stage of testis. Whereas in yak, the expression HSP70 reach top at adult and HSP90 at juvenile, different with cattle-yak, it’s HSP70 keep higher during every develop stage of testis (Fig. 3-III).

As shown in Fig 5, the protein expression levels of HSP70/90 were detected in the testicular tissues at different development stages of cattle-yak and yak. The results showed that the protein expression levels of HSP90 were significantly higher than those of HSP70 in almost all the developmental stages of cattle-yak (P<0.01). By contrast, the protein expression levels of HSP70 were significantly higher than those of HSP90 in almost all the developmental stages of yak (P<0.01), except in the newborn (Fig. 5-II, Fig. 6-III).

**Fig 5.**
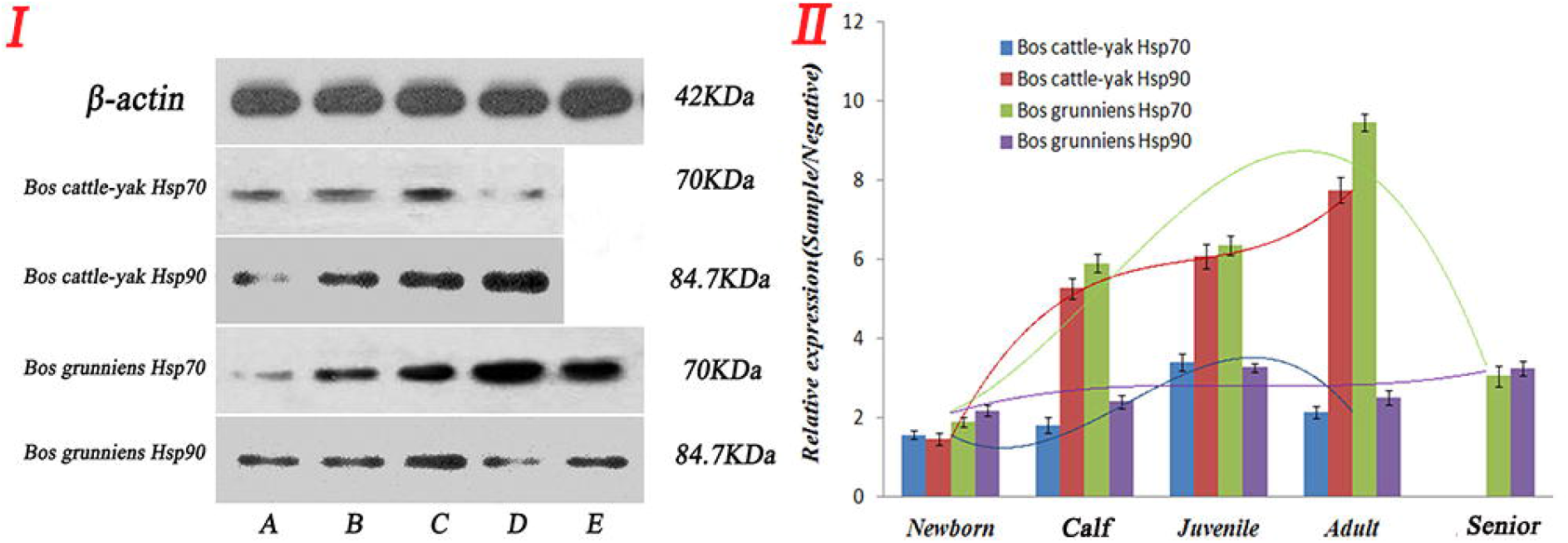
The protein expression in the testicular tissue of cattle-yak and yak. I. The schematic diagram labels bands and sizes. Different color represents expressions of HSP70/90 in the different testicular tissue of cattle-yak and yak.

The testicular seminiferous tubule of the newborn cattle-yak and yak was thin and sparse (Fig. 6-I and 6-II). As the cattle-yak aged, the number of spermatogenic cell at the different development stages decreased, and few sperm and sperm cells appeared. HSP90 was highly expressed in all levels of spermatocyte in the cattle-yak testicular tissue, whereas the expression levels in the basement-membrane and myogenic cells were the weakest. By contrast, the HSP70 expression levels in spermatocyte were the weakest. We are interested in what was showed, as the organism developed, seminiferous tubules became closely packed, luminals became enlargement, and different amounts of sperm cells appeared during the development of sperms, with the largest count occurring during the adult stage. For the elderly yak, the amount of spermatogenic cells in the seminiferous tubule decreased significantly. The HSP90 expression was strongest in the yak testis in primary spermatocyte, followed by that in the secondary spermatocyte. Sperm cells also had expression, contrary to the spermatogonium cells. The HSP70 expression levels in the mesenchymal cells were strong, while the expression levels in the basement-membrane and myogenic cells were the weakest.

**Fig 6.**
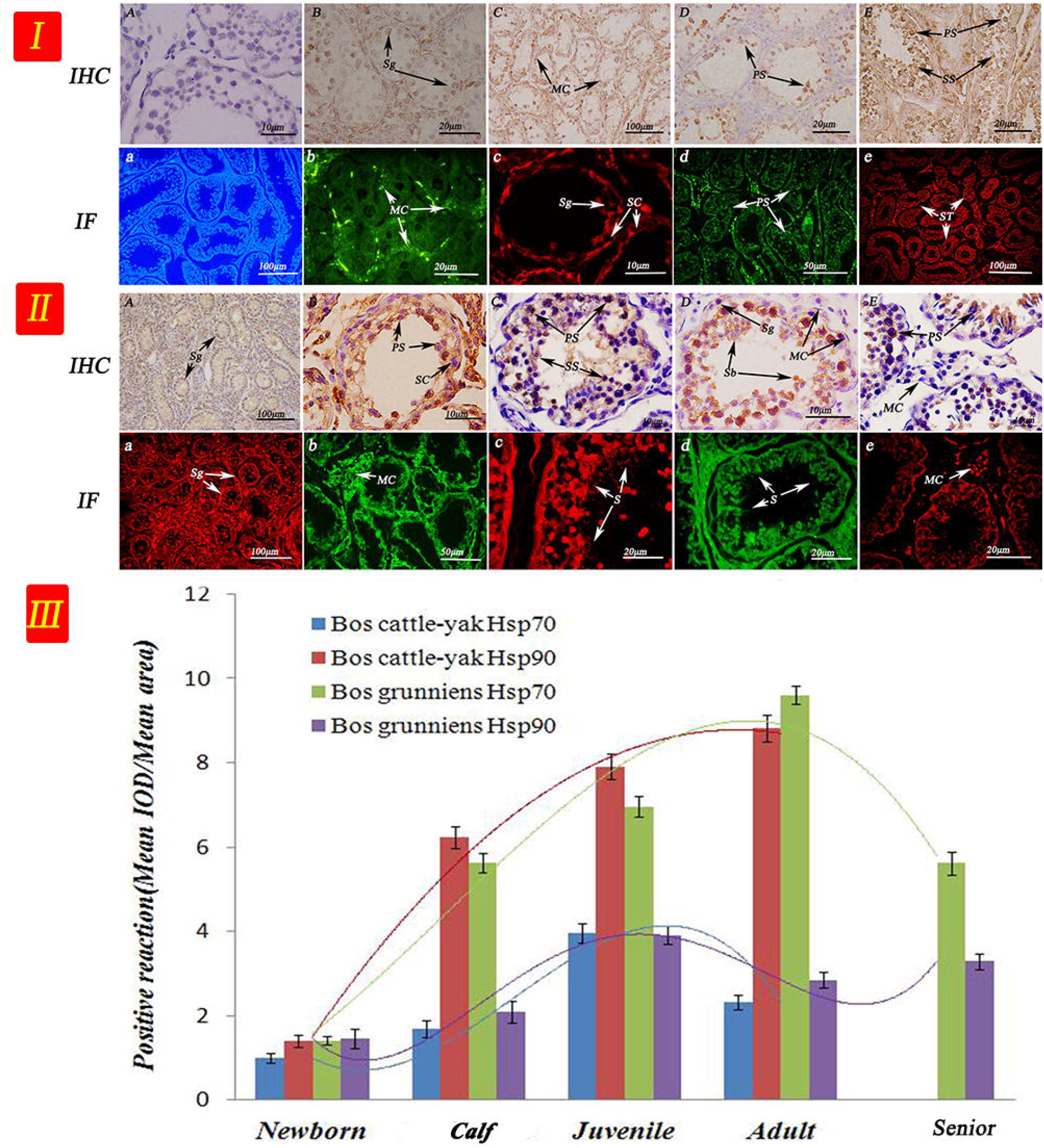
IHC and IF localization of HSP70/90 in the testicular tissue of cattle-yak and yak. (A) The control sections collected from the testicular tissue of adult yak, without immunoreactions (negative control). (B), (D) Positive staining for HSP70 was observed in the spermatogonium and primary spermatocyte of the newborn and juvenile testis. (C), (E) Positive staining for HSP90 was observed in mesenchymal, primary spermatocyte and secondary spermatocyte cells of the childhood and adult testis. (a) The control sections collected from the testicular tissue of adult yak, without immunoreactions (negative control). (b), (d) Positive staining for HSP90 was observed in the mesenchymal and primary spermatocyte of the newborn and adult testis. (c), (e) Positive staining for HSP70 was observed in the spermatogonium, sertoli cells and seminiferous tubule of the childhood and adult testis. (A), (C), (E) Positive staining for HSP90 was observed in the spermatogonium, secondary spermatocyte and mesenchymal cells of the newborn, juvenile and senile testis. (B), (D) Positive staining for HSP70 was observed in primary spermatocyte, sertoli cells, spermoblast and spermatogonium of the childhood and adult testis. (a),(c), (e) Positive staining for HSP70 was observed in the spermatogonium, sperm and mesenchymal cells of the newborn, juvenile and senile testis. (b), (d) Positive staining for HSP90 was observed in the mesenchymal cells and sperm of the childhood and adult testis. Note. Primary spermatocyte (PS), secondary spermatocyte (SS), spermatogonium (Sg), spermoblast (Sb), sperm (S),sertoli cells (SC), mesenchymal cells(MC), myoid cells (MC), seminiferous tubule (ST). The result of optical density analysis value. The curved lines represent the expression trend of HSP protein in different testis tissues, and different color represents different proteins.

## Discussion

This study is the first to isolate, sequence, and characterize cDNA clones that encode HSP90 from cattle-yak. Previous studies found that the protein space structure difference leads to function different of HSP90 (Agarwal et al. 2016). Using our reference sequence, our analytical results showed that three ORFs in the HSP90 sequence illustrated the frame-shifting property of the sequence. We also present evidence that susceptibility of trees to cattle-yak HSP90 associated with *yak, taurus, goats* were very high, and that cattle-yak evolution with the above animals had very close genetic relationship. In eukaryotic cells and the different HSP gene promoters upstream of TATA, heat shock element (HSE) with size of approximately 20 bp was also necessary in the transcription of specific nucleotide sequence (C-GAA-TTC-G) (Pelham and Bienz. 1982; Jenkins et al. 2016). The amino acid sequence of the various protein spatial structures differed. In humans, HSP90 vary according to their richness in glutamine (Yeyati and Heyningen. 2008). Results showed (Fig 2) that the cattle-yak amino acids happened multipoint mutation relative to yak. So, the molecular data show that HSP90 gene could be used as a single nucleotide polymorphism marker to evaluate the loss of genetic diversity caused by hatchery selection or internal control gene in cattle-yak. Thus, we believed that the amino acid mutation is an important factor for functional differences of HSP90.

The gene and protein expression of HSP70/90 was significant not only in the different species but also in the different tissues of the same species. Interestingly, compared with the males, the females had significantly lower post-exercise HSP72 levels which were not influenced by estradiol supplementation. This characteristic is mainly displayed in the heart, liver, lung and red and white vastus muscle (Paroo et al. 1999). In addition, HSPs under cold stress induced oxidative stress in spleen and influenced immune function in chicks (Zhao et al. 2014). It was hard to believe these data also verify the controversies regarding the appropriate physiological function of different tissues. We found that gene and protein expression of HSP70 was higher in the kidney, heart, and cerebellum of cattle. However, HSP90 had higher expression in the lung, cerebellum, and liver of cattle. Studies have shown that the apoptosis were reduced when HSP90 expression by inhibition (Hu et al. 2018). Luckily for us, considering the high expression in the cerebellum and heart of cattle-yak and yak, we speculated that HSP70/90 could protect cells and tissue from the environmental stimulus.

Previous studies demonstrated that HSP90 can protect lung tissue from damage caused by cold air. Different stimuli could induce HSPs in the proximal with significant differential expression in the bronchioles and respiratory bronchiole epithelial cells and smooth muscle cells (Manjili et al. 2002; Li et al. 2015). Gene and protein expression levels of HSP70/90 were significantly upregulated when the porcine fetal fibroblast and oviduct epithelial cells were stimulated by cold (LI et al. 2012). Studies have shown that HSP70/90 played a regulatory role in the adaptation of japonicus to high temperature and low salinity (Wang et al. 2014). This study demonstrated the localization of HSP70/90 in the cell membrane and cytoplasm, not in the nuclei, of organs. It is undeniable that, HSP70/90 plays an active function in the tissue physiological adaptation in cattle-yak and yak.

We performed studies to examine which expression was altered by gene and protein in cattle-yak to elucidate the mechanism by which HSP70/90 affect the development of testicular tissue. Encouragingly, HSP90 expression was generally higher than that of HSP70 in the cattle-yak testicular tissue, especially during childhood, juvenile, and adulthood. Spinaci reported that the expression level of HSP90 was gradually reduced when the pig sperm was frozen, and the protein content was reduced before sperm quality declined (Spinaci et al. 2016). Meanwhile, the expression in the cattle-yak remained at a higher level from juvenile and adulthood. This result is consistent with the results obtained by Abd El-Fatta, who showed that the high expression of HSP90 could inhibit testicular dysfunction caused by DEHP (El-Fattah et al. 2016). For the male cattle-yak infertility, the expression of HSFY2 was delayed, not absent. The fertility of cattle-yak was restored, as the HSFY2 was expressed (Chai et al. 2014; Sarge and Cullen. 1997). In addition, the HSP70/90 lower expression led sperm quality to low of boar (Zannoni et al. 2017). Hsp70 expression may have been upregulated as a protective mechanism against apoptosis in spermatozoa of infertile men (Erata et al. 2008). Thus, as the Fig 7, we speculated that the high expression of HSP90 in the testicular tissue of male cattle activate offspring male cattle-yak to restore normal fertility.

**Fig 7.**
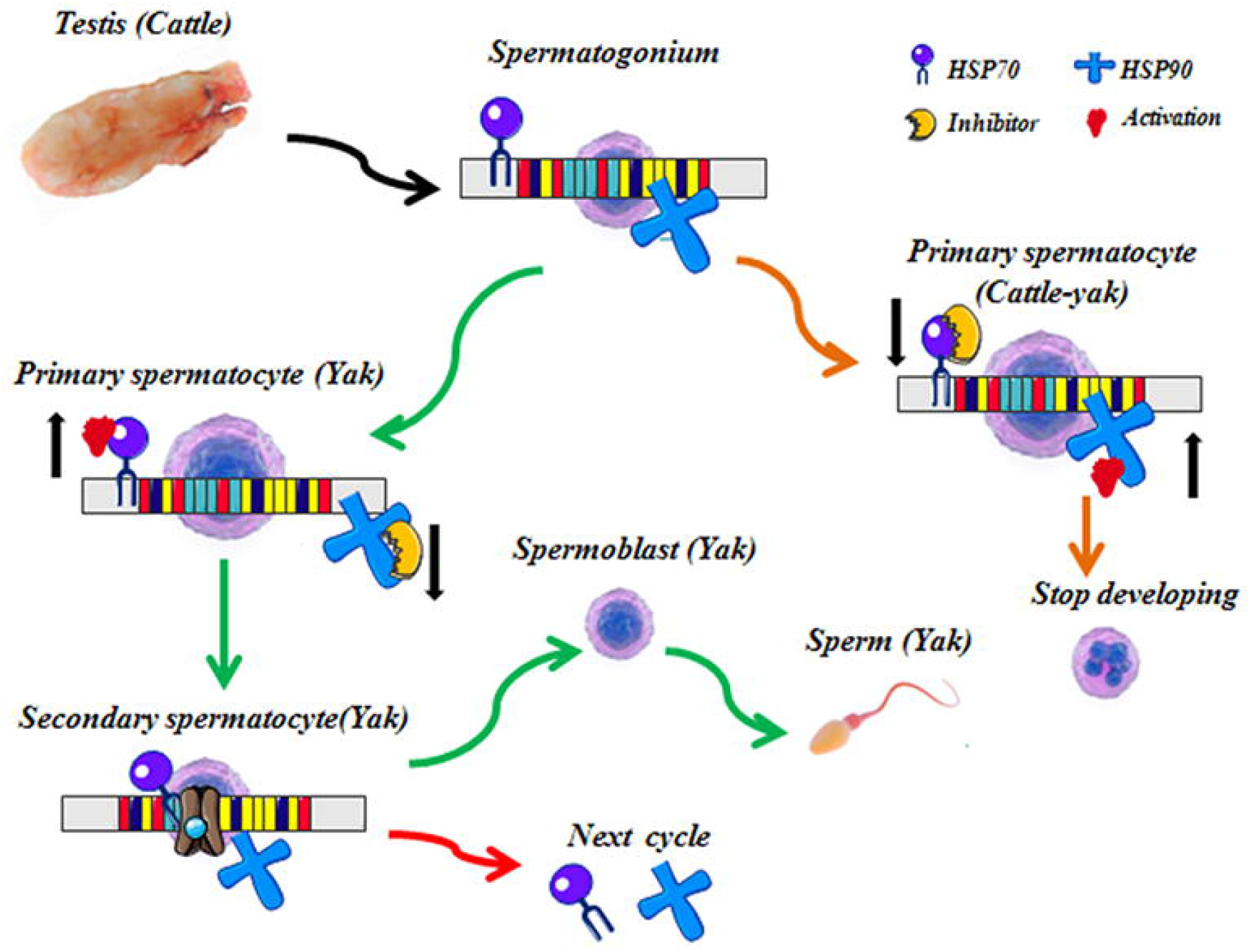
The HSP70/90 schematic diagram of action mechanism in the testicular tissue. The higher expressed HSP90 could block the development of the primary spermatocyte in testis of cattle-yak. The higher expressed HSP70 and the decreased HSP90 could promote the sustrained development from the spermatocyte to mature sperm in testis of yak.

We analyzed the HSP70/90 expression in the testicular tissue during the different stages of yak development to establish whether the HSP70/90 content has positive correlations with the reproductive capacity of yak. The results indicated that the highest HSP70-2 gene expression level was in the yak testicular tissue, followed by those of the cattle and cattle-yak (Yl et al. 2016). In addition, HSP 70 could protect against sperm apoptosis when received the stimulation by high temperature (Mann et al. 2002). This result is largely consistent with our experimental result that the HSP70 expression level in the testicular tissue of yak increased gradually with age, and the strongest expression was during the adult stage, then gradually decreased in old age. Dix reported that mutant HSP70-2 could inhibit the coupling of nuclear chromosome coming into meiosis (Dix et al. 1996). Furthermore, the HSP90 firstly increases and then decrease at testicular development process of rabbit (Wu et al. 2011). What makes this amusing, as the Fig 7, we speculated that the low expression of HSP90 and high expression of HSP70 were activated the normal reproduction of yak.

The immune response of the HSP family protein was mainly concentrated in the epithelial tissue and cell serous. Cedenho found that HSP70-2 was expressed in the normal testis and testicular tissue with disturbance of sperm maturation (Cedenho et al. 2006). However, the expression was higher in the normal testicular tissue and spermatoblast serous, lower in the testicular tissue and spermatoblast serous with disturbance of sperm maturation. No expression was found in the testicular tissue of sustentacular cell syndrome patients. Economou proved stage and cell-specific expression and intracellular localization of the small Hsp27 during oogenesis and spermatogenesis in the Mediterranean fruit fly (Economou et al. 2017). The expression was strongest in the yak testis in the primary spermatocyte, followed by the secondary spermatocyte. Sperm cells also showed expression, contrary to spermatogonium cells. Thus, we demonstrated that the low expression of HSP70 was related to male sterility. HSP90 was highly expressed in all levels of spermatocyte of cattle-yak testicular tissue, especially in the mesenchymal cells. In 2003, Cardozo declared the evident increase in the intensity of immune response in the testicular tissue with disturbance of sperm production, with the most obvious effect in the myoid cells (Cardozo et al. 2003). In addition, the lack of sperm caused infertility and hastened spermatocyte apoptosis in the spermatogenic epithelium. Even more remarkably, the mule, a classic example of hybrid sterility in mammals also exhibits a similar spermatogenesis breakdown, it failure at the first meiotic stage (Steiner and Ryder. 2013). Thus, the high expression of HSP90 caused the blockage of spermatocyte development. Is most proud of is, this study showed an important breakthrough in the study of male infertility.

## Conclusions

The following conclusions can be drawn from this study. The amino acids were changed that the main cause for protein functional differences of HSP90. HSP70/90 has an obvious differential expression in different organs and differences age paragraph testicles of cattle-yak and yak. In addition, HSP70/90 was mainly detected in epithelial of tissue, spermatogenic cell and mesenchymal in the testicular. At the core, HSP70/90 showed the specific expression in yak and cattle-yak testicular tissue, and it has important correlation with cattle-yak infertility

## Materials and Methods

### 1. Experimental animals

Cattle were sampled from the pasture in the Tibetan plateau in Qinghai, China (Gu et al. 2015). The study was approved by the Animal Ethics Committee of Gansu Agricultural University. All experiments were performed in accordance with relevant guidelines and regulations. A total of 54 healthy cattle (24 cattle-yak and 30 yaks) were included in this study (Table 1), All cattle were kept under the same natural conditions (altitude, approximately 2,300 m; temperature, 2–5 °C; and oxygen content, 14.97%) (Wiener et al. 2003). Male cattle-yak without feeding value for herders were excluded, so no samples for the senile group were included. The cattle with conventional disease were considered healthy individuals.

### 2. Preparation of mRNA and proteins

Tissue samples were selected from the healthy cattle subjects. A total of 21 tissues were dissected, collected in frozen liquid nitrogen, and then stored at -80 °C before use. The RNA from the cattle tissues were isolated using a TRIzol kit (R1100, USA), and reverse-transcription polymerase chain reaction (RT-PCR) was conducted to clone the cDNA.

Frozen samples were ground in liquid nitrogen and transferred to centrifuge tubes with RIPA/PMSF (Solarbio, China). After fully blending and mixing until a pink color was obtained, the sample tubes were incubated on a spiral oscillator for 2 h (200 r/h) on ice. After centrifuging at 4 °C for 10 min (12000 r/h), the complete divided protein was obtained. In addition, the total protein concentration for each sample was measured.

The total protein concentration was then adjusted to the same level, and 4× sample buffer at 100 °C was added for 12 mins to completely denature the proteins.

### 3. Molecular cloning of HSP90

The degenerate primers used to amplify the HSP90 sequence were based on the published partial sequence of HSP90 mRNA in the National Center for Biotechnology Information (NCBI) database (Accession No. NM-001012670.2). Primers for cloning the initial fragment of HSP90 mRNA were designed according to the predicted conserved sequences in other *Bos* cattle-yak animals (Table 2). The amplified segments were inserted into the cloning vector pMD-18T and were transfected into *Escherichia coli* JM109 competent cells. The primers for 5‘ HSP90 and 3‘ HSP90 were designed using the sequencing data. Then, the segments from 5‘ to 3‘ HSP90 from the first-strand cDNA were used for cloning and sequencing.

### 4. Analysis of HSP90 protein

The open reading frames (ORF) in the complete mRNA sequence of HSP90 were identified using an ORF finder (http://www.ncbi.nlm.nih.gov/projects/gorf/orfig.cgi), and then the nucleotide sequences were translated into amino acids using the Vector NTI 11 software (He et al. 2016). The HSP90 sequence was analyzed using the ODC HSP90 finder software, and the codon region and frameshifting site were identified (Gandre et al. 2003).

Homology searches were performed using BLASTn and BLASTp in NCBI. The Conserved Domain (CD) Search service was used to identify the CDs in the predicted protein sequences (http://www.ncbi.nlm.nih.gov/Structure/cdd/cdd.shtml). The 3-D structure of the predicted protein was predicted according to the methods described in the website (http://bioinf.cs.ucl.ac.uk/psipred/). The deduced amino acid sequence of HSP90 was aligned using the CLC main workbench software (http://www.clcbio.com) with known homologous proteins of the HSP90 class obtained from the GenBank. A phylogenetic tree was constructed using the CLC main workbench software through the neighbor-joining method for the amino acid sequences of HSP90 from the SwissProt databank/Genbank.

### 5. Expression of HSP70/90 gene in the different tissues

The distribution of HSP70/90 in the tissues of cattle-yak and yaks were detected during the development of testicles and in the non-reproductive system (kidney, heart, cerebellum, liver, lung, and spleen). The expression levels of HSP70/90 in the different tissues were detected using quantitative real-time PCR (Invitrogen, USA) with HSP70/90 specific primers. Actin was used as a reference gene to normalize the amount and quality of each cDNA, because this gene is expressed constitutively in the different tissues (Gandre et al. 2003).

### 6. Protein expression of HSP70/90 in testicle tissues

Protein samples was thawed, mixed and separated into beads using a spin column (Bio-Rad), and then separated on a 5% SDS-PAGE gel for Western blot analysis. After electrophoresis, the proteins were transferred from the gel onto NC membranes (Millipore Corporation, Billerica, MA, USA). The membranes containing protein were blocked with 5% fat-free milk in TBST at room temperature for 2 h and hybridized using HSP70/90 Abcam (1:1000) or rabbit polyclonal Abcam (1:2000) at 4 °C overnight. The membrane was then washed four times with TBST and labeled with HRP-conjugated secondary Ab (1:4000) for 2 h at room temperature. After washing five times with 1× TBST, HSP70/90 was detected on the membrane with an ECL detection kit (Beyotime, China). The intensities of the bands on the blots were measured using a densitometric analysis system (Bio-Rad). The intensities of the β-actin bands were used for normalization (Hu et al. 2016).

### 7. Immunofluorescence and Immunohistochemical Assays

The main organs and tissues of the cattle-yak and yak were fixed in 4% paraformaldehyde solution at room temperature for hebdomad. Tissue pieces were clipped and paraffin-embedded, and the sections were sliced, dried, and stored.

Samples were dewaxed using dimethyl benzene and then dehydrated with an increasing alcohol gradient for immunohistochemical staining to investigate the HSP70 and HSP90 expression levels. The endogenous peroxidase was eliminated with 3% deionized H_2_O_2_ (18–22 min), and the sections were rehydrated and sealed with goat serum (18–22 min). After overnight incubation at 4 °C with the primary rabbit anti-HSP70 and mouse anti-HSP90 monoclonal antibodies (1:300, Abcam, Hong Kong), the sections were then incubated with the secondary antibody. Then, the labeled samples were counterstained with 3,3‘-diaminobenzidine, and the nuclei were appeared (Gu et al. 2015). In addition, two antibodies showed different colors in immunofluorescence, and redyeing the nuclei was not necessary.

### 8. Measurement and statistical analyses

The intensity in the Western blot images and immunofluorescence and immunohistochemical assays were measured using integrated optical density and Image-Pro plus 6.0. Data were analyzed using SPSS 21.0. Spearman correlation of coefficients was analyzed between β-actin and sample protein levels. Other data were analyzed by one-way ANOVA and Duncan‘s post hoc test. P-values less than 0.05 between groups were considered statistically significant (He et al. 2016).

## Acknowledgements

Yan Cui and Penggang Liu designed the experiments, performed the experiments, analyzed and interpreted data, and wrote and edited the manuscript.

Jun Liu, Liangli Song, Yuanfang Xu and Shengnan Zou performed the experiments, analyzed and interpreted data for manuscript.

Sijiu Yu, Junfeng He, Qian Zhang and Ting Wang contributed reagents/materials/analysis tools for manuscript.

